# Nerve Growth Factor Signaling Tunes Axon Maintenance Protein Abundance and Kinetics of Wallerian Degeneration

**DOI:** 10.1101/2024.12.31.630780

**Authors:** Joseph A. Danos, Merve Addemir, Lily McGettigan, Daniel W. Summers

**Affiliations:** Department of Biology, University of Iowa, Iowa City, IA 52242 USA; Iowa Neuroscience Institute, University of Iowa, Iowa City, IA 52242 USA

## Abstract

Neurotrophic factors are critical for establishing functional connectivity in the nervous system and sustaining neuronal survival through adulthood. As the first neurotrophic factor purified, nerve growth factor (NGF) is extensively studied for its prolific role in axon outgrowth, pruning, and survival. Applying NGF to diseased neuronal tissue is an exciting therapeutic option and understanding how NGF regulates local axon susceptibility to pathological degeneration is critical for exploiting its full potential. Our study identifies surprising connections between NGF signaling and proteostasis of axon maintenance factors. NGF deprivation increases Nmnat2 and Stmn2 protein levels in axon segments with a corresponding delay in Wallerian degeneration. Conversely, acute NGF stimulation reduces local abundance of these axon maintenance factors and accelerates Wallerian degeneration. Pharmacological studies implicate phospholipase C as the key effector in TrkA activation, which drives degradation of palmitoylated Stmn2. While seemingly opposed to neuroprotective activities well-documented for NGF, downregulating Nmnat2 and Stmn2 favors axonal outgrowth over transient hyper-susceptibility to Sarm1-dependent degeneration. This new facet of NGF biology has important implications for axonal remodeling during development and sustained integrity through adulthood.

## INTRODUCTION

Neurons extend long axons necessary for functional communication through the nervous system. Axons can reach over a meter in length in some contexts and are uniquely vulnerable to stressors that occur during aging, physical trauma, and metabolic stress. Axon degeneration is a common event in a wide variety of neurodegenerative disorders and boosting axonal resilience has broad therapeutic potential (Coleman and Hoke, 2020). Identifying factors regulating axon susceptibility to pathological degeneration offers great value toward fulfilling this goal.

Considerable insight on pathological axon degeneration comes from models of Wallerian Degeneration in which axotomy triggers dismantling and fragmentation of disconnected distal axons (Wang *et al*., 2012; Coleman and Hoke, 2020). Axotomy deprives distal axons of short-lived maintenance factors such as Nmnat2 and Stmn2 (Gilley and Coleman, 2010; Shin *et al*., 2012). Nmnat2 depletion stimulates Sarm1 NAD^+^ activity and a cascade of self-destructive events including cytoskeletal dismantling and phosphatidylserine exposure, culminating in loss of membrane permeability and axon fragmentation (Gilley *et al*., 2015; Figley and DiAntonio, 2020; Ko *et al*., 2021). Either elevating Nmnat2 or inhibiting Sarm1 prolongs functional survival in pre-clinical models of neurodegeneration reinforcing therapeutic potential of this pathway (Krauss *et al*., 2020; Arthur-Farraj and Coleman, 2021; Geisler, 2024).

In contrast to pathological axon destruction, the pruning of excess axonal processes is critical for establishing functional neuronal circuits during development (Luo and O’Leary, 2005; Saxena and Caroni, 2007). Key to successful innervation in sympathetic and sensory systems is nerve growth factor (NGF) binding to tropomyosin related kinase A (TrkA) and stimulation of a PI3K-mediated pro-survival retrograde signal to the neuronal soma (Yao and Cooper, 1995). NGF deprivation mobilizes a DLK-MKK4/7-JNK signaling complex that induces caspase-dependent cell death and axon degeneration (Sengupta Ghosh *et al*., 2011; Holland *et al*., 2016; Simon *et al*., 2016; Niu *et al*., 2022). Supplementing NGF shows considerable therapeutic promise in preventing retinal degeneration and slowing Alzheimer’s disease (Lambiase *et al*., 2009; Amadoro *et al*., 2021). However, systemic NGF application causes hyperalgesia (Lewin *et al*., 1993; Petty *et al*., 1994) and anti-NGF treatments are utilized in pain management (Wise *et al*., 2021). Mechanistic underpinnings of these seemingly contradictory responses to NGF stimulation are not clear.

Developmental axon pruning and pathological axon degeneration are often treated as separate pathways operating at distinct stages in an organism’s lifespan. There are notable points of convergence suggesting potential for cross-regulation. Death receptor 6 promotes axon degeneration in response to NGF deprivation and axotomy (Gamage *et al*., 2017). Calpain proteases promote dismantling of neurofilaments downstream of caspase proteases and Sarm1 in both contexts (Yang *et al*., 2013; Ko *et al*., 2021). Activating DLK-MKK4/7-JNK accelerates degradation of Nmnat2 thereby hypersensitizing axons to Sarm1-dependent degeneration (Summers *et al*., 2020). In this study we identify a surprising connection between NGF signaling and proteostasis of axon maintenance factors. NGF deprivation increased local Nmnat2 and Stmn2 abundance with a corresponding delay in fragmentation of severed axons. Conversely, acute NGF stimulation reduced levels of these palmitoylated axon maintenance factors and accelerated Sarm1-dependent degeneration. Our results point to unexpected influence for local NGF signaling on axon vulnerability through regulated degradation of axon maintenance factors.

## RESULTS

### Blocking NGF signaling through TrkA delays Wallerian Degeneration

NGF deprivation stimulates a retrograde DLK-MKK4/7-JNK signaling complex responsible for triggering apoptotic cell death (Sengupta Ghosh *et al*., 2011). Since activating this MAPK pathway enhances Nmnat2 degradation and accelerates Wallerian Degeneration we predicted acute NGF deprivation would likewise accelerate fragmentation of severed axons. To test this prediction, we removed NGF from mouse, embryonic-derived Dorsal Root Ganglia (DRG) sensory neurons four hours prior to axotomy with a razor blade. Fresh media lacking NGF was supplemented with anti-NGF antisera to inactivate residual NGF protein (Levi-Montalcini and Booker, 1960). Media containing NGF was exchanged on control cells to account for this manipulation in our experiments. Severed axons were visualized with an automated microscope once an hour over a twelve-hour period and axon degeneration quantified with an ImageJ macro that calculates fragmented axons in a field based on object circulatory (Gerdts *et al*., 2011). In the presence of NGF there was a lag phase of approximately four hours in which no change in axon morphology occured. Axon fragmentation ensued after this lag phase and plateaued as the entire axon field degenerated. Contrary to our prediction, NGF deprivation delayed the onset of axon fragmentation and complete axon degeneration was not reached during the experimental timecourse (Figure 1A).

**FIGURE 1.**
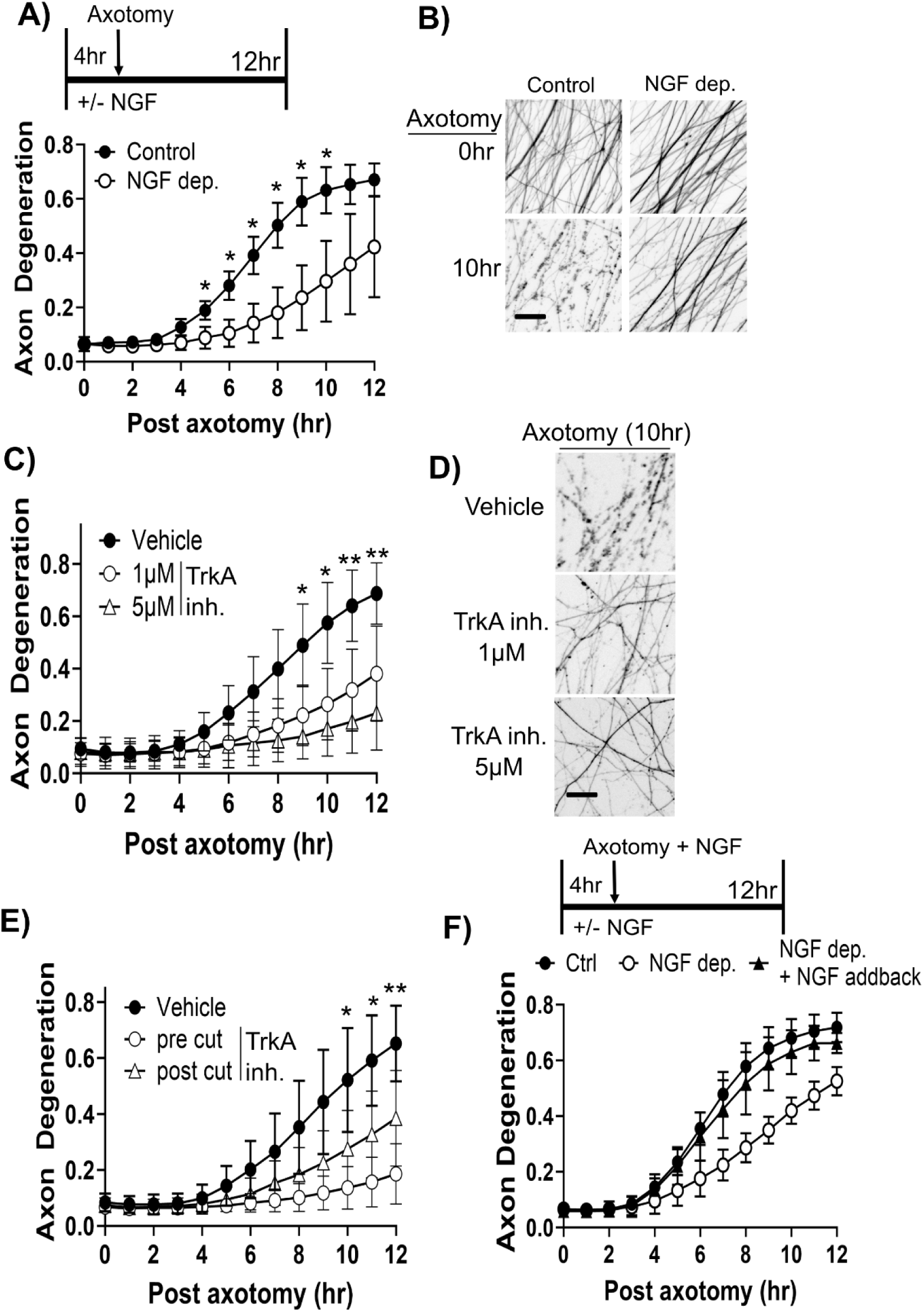
NGF deprivation delays fragmentation of severed axons. **(A)** DRG sensory neurons were cultured in media with or without NGF for four hours prior to manual axotomy with a razor blade. In NGF minus conditions media also contained anti-NGF antisera. Axon degeneration was measured from severed axons each hour for a twelve-hour period (N=3). **(B)** Representative images of severed axons at 10hr post axotomy. **(C)** Neurons were pretreated with two different doses of a TrkA inhibitor (GW441756) four hours prior to axotomy. Example images are shown in **(D)** (N=4, asterisks refer to statistical comparisons between Vehicle and 5 µM dose). **(E)** GW441756 (5µM) was applied four hours prior to axotomy (pre cut) or immediately after axotomy (post cut) (N=4, asterisks refer to statistical comparisons between Vehicle and pre cut). **(F)** NGF deprivation was performed as in **(A)** except anti-NGF antisera was omitted. In addback condition NGF was applied immediately post-axotomy. Statistical comparisons in timelapse experiment performed with a Repeated Measure Two-way ANOVA *p<0.05, **p<.01 (N=4). Scale bar = 20 µm. Error bars represent +/-1 STD.

As a complementary approach sensory neurons in media containing NGF were treated with a small molecule TrkA inhibitor (GW447156). Consistent with our findings using acute NGF deprivation, TrkA inhibition four hours prior to axotomy delayed axon degeneration in a dose-dependent manner (Figure 1C & D). The ease of this pharmacological approach inspired us to evaluate whether TrkA inhibition post-axotomy was sufficient to delay axon degeneration. Applying TrkA inhibitor immediately after axotomy delayed axon degeneration albeit to a diminished extent compared to pre-cut treatment (Figure 1E). Therefore, local NGF deprivation delays Wallerian degeneration however this effect is most potent after prolonged deprivation in intact axons.

We next evaluated whether reapplying NGF after axotomy would restore kinetics of axon degeneration. To conduct this experiment NGF deprivation was performed without anti-NGF antisera to enable NGF reapplication. NGF deprivation delayed axon degeneration in this experiment though not to the same extent as observed in the presence of anti-NGF antisera. Reapplying NGF immediately following axotomy restored kinetics of axon degeneration to a similar rate as observed in controls containing NGF (Figure 1F). Therefore, local NGF signaling affects the rate of fragmentation in severed axons.

### NGF deprivation increases Nmnat2 and Stmn2 abundance in axon segments

Elevating Nmnat2 suppresses Sarm1 activation and extends survival of severed axons (Gilley *et al*., 2015; Figley *et al*., 2021). We predicted NGF deprivation increases Nmnat2 protein. Axon-only extracts were collected from sensory neurons undergoing NGF deprivation for four hours then evaluated by western immunoblotting. NGF deprivation increased endogenous Nmnat2 protein approximately 2-fold (Figure 2A). Another axon maintenance factor called Stmn2 is frequently co-regulated with Nmnat2 in axons (Summers *et al*., 2018; Summers *et al*., 2020). Moreover, Stmn2 protein levels are reduced in motor neurons with TDP-43 cytoplasmic aggregates (Klim *et al*., 2019; Melamed *et al*., 2019). We detect a 2-fold increase in Stmn2 protein from axon-only extracts after NGF deprivation. Applying a TrkA inhibitor for two hours likewise increased Stmn2 and Nmnat2 protein levels 1.5-fold increase over vehicle control in axon-only extracts (Figure 2B). As a separate approach, we measured fluorescence intensity from exogenously expressed Stmn2-Venus. TrkA inhibition increased Stmn2-Venus fluorescence intensity 1.4-fold in axon segments (Figure 2C).

**FIGURE 2.**
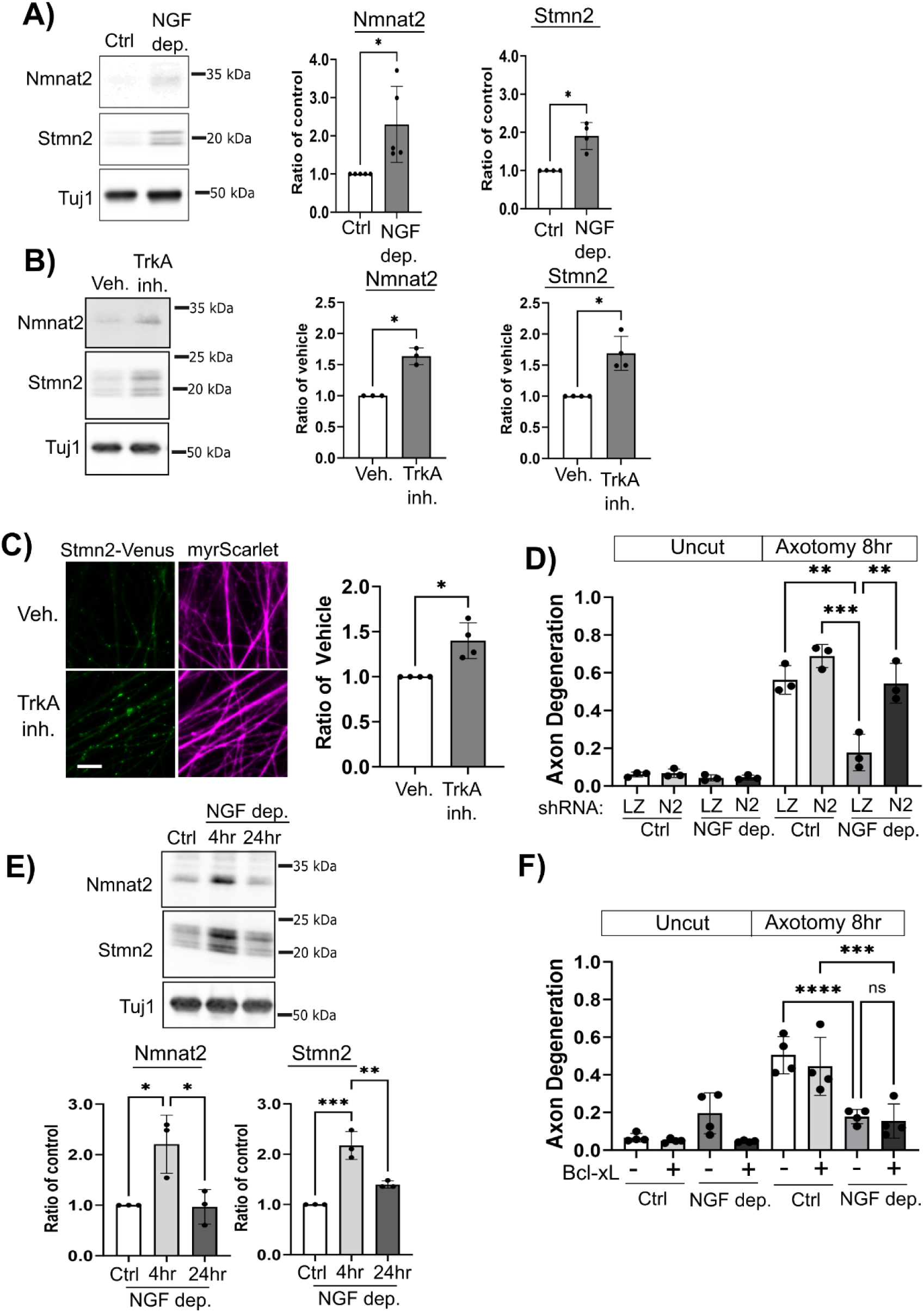
Acute NGF deprivation increases axonal Nmnat2 and Stmn2 protein. **(A)** Western blots of Nmnat2 and Stmn2 from axon-only extracts collected from control or NGF-deprived neurons (4hr) with quantification on the right for Nmnat2 (N=5) and Stmn2 (N=4). **(B)** Western blots of Nmnat2 and Stmn2 from axon-only extracts collected from vehicle or GW441756 (5 µM) treated neurons (4hr) with quantification on the right for Nmnat2 (N=3) and Stmn2 (N=4). **(C)** Images of distal axons from Stmn2-Venus expressing neurons were treated with vehicle or GW441756 (5 µM) for four hours with quantification on the right (N=4). **(D)** NGF deprivation was performed as described in Figure 1A on neurons transduced with lentivirus expressing shLacZ control (LZ) or shNmnat2 (N2). Axon degeneration was measured eight hours post axotomy. **(E)** Nmnat2 and Stmn2 protein levels return to baseline in axon-only extracts after 24hr NGF deprivation. Quantification is shown on the right (N=3). **(F)** Neurons were transduced with lentivirus expressing Bcl-xL or an empty vector lentivirus. NGF deprivation was performed as described in Figure1A and axon degeneration measured eight hours after severing with a razor blade. For A-D, statistical comparisons performed with Welch’s t-test. In D & F, one-way ANOVA with post-hoc unpaired t-tests were performed. For all statistical tests *p<0.05, **p<0.01, and ***p<0.005. Error bars represent +/-1 STD.

If elevated Nmnat2 is responsible for extending survival of severed axons then reducing Nmnat2 should suppress axon protection afforded by NGF deprivation. To test this prediction, we introduced an shRNA targeting Nmnat2 via lentiviral transduction, performed NGF deprivation, and measured degeneration of cut axons. Since prolonged Nmnat2 depletion can spontaneously induce Sarm1-dependent axon degeneration we controlled the timing of shRNA application such that uncut axons were intact during the experimental period. NGF deprivation suppressed axon degeneration ten hours post axotomy in the presence of a control shRNA (shLacZ). However, knocking down Nmnat2 reversed axon protection during NGF deprivation indicating this maintenance protein is required for extended axon survival in this model (Figure 2D).

We next evaluated whether Nmnat2 and Stmn2 protein levels remain elevated during prolonged NGF deprivation. Embryonic-derived sensory neurons undergo caspase-dependent cell death and axon degeneration in response to extended NGF deprivation. To circumvent this restriction, we constitutively expressed the anti-apoptotic protein Bcl-xL to suppress caspase activation and prolong neuron survival in the absence of NGF (Garcia *et al*., 1992). Importantly, Bcl-xL overexpression does not block Sarm1-dependent Wallerian degeneration (Vohra *et al*., 2010)(Supplementary Figure 1). After twenty-four hours in media lacking NGF, Nmnat2 and Stmn2 protein returned to levels observed under normal NGF conditions (Figure 2E). Bcl-xL overexpression did not affect axon protection afforded by NGF deprivation after axotomy suggesting caspase activation is not required for this effect (Figure 2F).

### Acute NGF stimulation decreases axonal levels of Nmnat2 and Stmn2

If transient NGF deprivation boosts axonal Nmnat2 and Stmn2 protein, we predicted acute NGF stimulation would do the opposite and reduce protein levels. To model acute NGF stimulation we cultured DRG sensory neurons in the presence of NGF until Days *in vitro* (DIV) 6 then exchanged media lacking NGF for twenty-four hours. We transduced neurons with lentivirus overexpressing Bcl-xL to suppress caspase activation and prevent apoptotic cell death. On DIV7 we applied NGF to these cultures for two hours and measured endogenous Nmnat2 and Stmn2 protein from axon-only extracts. Nmnat2 and Stmn2 protein levels decreased 60% and 50% respectively after NGF application (Figure 3A). We also visualized endogenous Stmn2 in axon segments by immunofluorescence and detected a 37% decrease after NGF application (Figure 3B).

**FIGURE 3.**
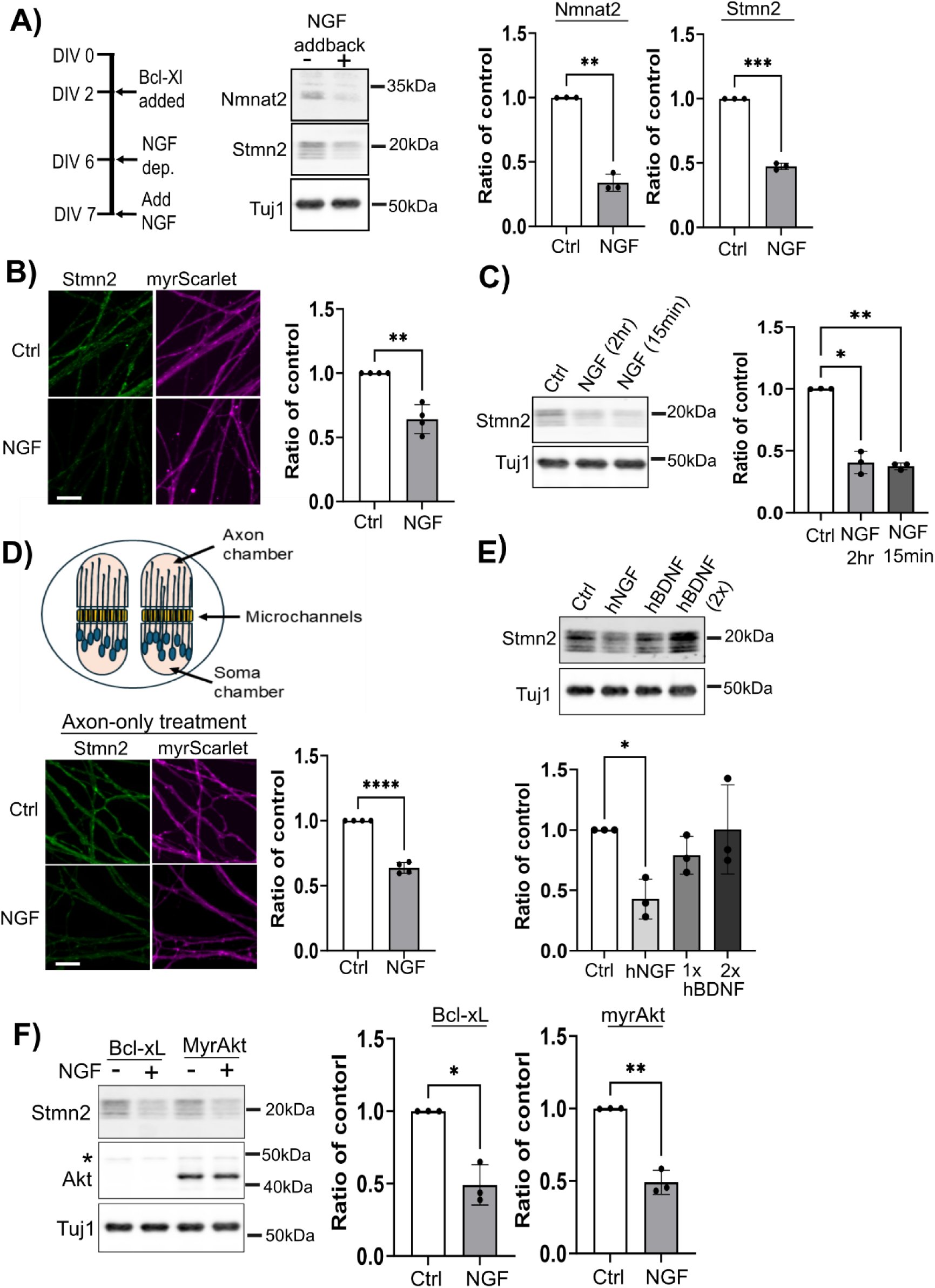
Acute NGF stimulation reduces axonal Nmnat2 and Stmn2 protein. **(A)** DRG sensory neurons underwent NGF deprivation for twenty-four hours prior to NGF application. Lentivirus expressing the anti-apoptotic protein Bcl-xL maintained neuron survival during the experimental period. NGF application for two hours reduced Nmnat2 and Stmn2 protein levels in axon-only extracts. Quantification is shown on the right (N=3). **(B)** Immunofluorescence of endogenous Stmn2 from DRG neurons after two-hour treatment with NGF. Quantification is shown on the right (N=4) **(C)** NGF was applied for fifteen minutes, replaced with media lacking NGF, then axon-only extracts collected two hours post treatment. Quantification is shown below (N=3). **(D)** DRG neurons were cultured in microfluidic devices to enable axon-only treatment with NGF for two hours. Quantification of Stmn2 immunofluorescence in the axon chamber is shown on the right (N=4). **(E)** Human BDNF (hBDNF) or human NGF (hNGF) were applied to NGF-deprived neurons for two hours where 1x and 2x dosages refer to 50ng/mL and 100ng/mL respectively. Western blots are from axon-only extracts with quantification below (N=3). **(F)** Overexpression of constitutively active AKT did not affect steady state Stmn2 levels in axons or NGF-induced Stmn2 loss. Quantification is shown on the right (N=3). The asterisk identifies endogenous AKT migrating slower than the truncated, constitutively active form. All statistical comparisons were performed with Welch’s t-test where *p<0.05, **p<0.01, ***p<0.05, and ****p<0.001. Error bars represent +/-1 STD. Scale bar = 10µm.

NGF binding stimulates TrkA endocytosis where this activated receptor can stimulate pro-survival signaling on an endosome (Yamashita and Kuruvilla, 2016). Accordingly, transient NGF exposure should be sufficient to provoke Stmn2 reduction. We applied NGF for fifteen minutes, washed neurons with media lacking NGF, and collected axon-only extracts two hours later. Fifteen-minute NGF treatment reduced Stmn2 protein levels to a similar extent observed after two-hour NGF treatment (Figure 3C).

We next employed microfluidic chambers to evaluate whether NGF stimulation selectively in the axon compartment is sufficient to decrease Stmn2 protein. Primary DRG sensory neurons were seeded in microfluidic chambers then subjected to the NGF withdrawal and addback paradigm described above. Cells were fixed and endogenous Stmn2 protein in the axon chamber detected by immunofluorescence. Applying NGF to the axon compartment for two hours decreased endogenous Stmn2 protein by 35% (Figure 3D) supporting the role of local NGF signaling in regulating Stmn2 abundance.

Control experiments were performed to address the specificity of NGF-induced Stmn2 depletion. We applied brain-derived neurotrophic factor (BDNF) to NGF-deprived neurons which can sustain neuron survival through TrkB (Deppmann *et al*., 2008; de Leon *et al*., 2021). Recombinant human BDNF was used in this experiment. As a species-specific control, we applied recombinant human NGF (hNGF) and observed a 55% decrease in Stmn2 protein from axon-only extracts (Figure 3E) similar to mouse NGF used in earlier experiments. However, human BDNF did not elicit an effect on Stmn2 protein levels even when applied at double the concentration of hNGF.

Bcl-xL overexpression functions at the level of mitochondrial cytochrome c release to suppress caspase activation. As an alternatively strategy to sustain survival signaling, we overexpressed a membrane-tethered, truncated form of Akt lacking its autoinhibitory Pleckstrin Homology domain (Kohn *et al*., 1996). NGF deprivation and addback were performed as described above. Constitutively active Akt did not alter baseline Stmn2 protein levels from axon-only extracts. NGF application in the presence of constitutively active Akt reduced Stmn2 protein to comparable levels observed with Bcl-xL overexpression. (Fig. 3F). Altogether, acute NGF stimulation decreases Stmn2 and Nmnat2 protein levels in axon segments.

### TrkA activation is responsible for NGF-induced Stmn2 reduction

NGF signaling through the high-affinity TrkA receptor is well-studied for roles in axon outgrowth and neuron survival (Kaplan and Stephens, 1994). However, cooperation with the low affinity receptor p75 also regulates NGF-TrkA signaling (Hempstead *et al*., 1991). Signaling through the p75 receptor is more closely linked to neurodegeneration which would be consistent with our observation (Khan and Smith, 2015; Meeker and Williams, 2015). We employed the NGF addback paradigm described in Figure 3 in combination with pharmacology and genetic manipulation to determine whether NGF reduces Stmn2 protein through the TrkA or p75 receptor. We used Stmn2 protein levels as our primary readout in most of our subsequent experiments because reagents for detecting this microtubule-binding protein are reliable and well-validated.

Co-applying NGF with a TrkA inhibitor suppressed NGF-induced reduction in Stmn2 protein levels (Figure 4A). Conversely, CRISPR-editing of the p75 gene did not affect NGF-induced reduction in Stmn2 protein (Figure 4B) though endogenous p75 protein levels were substantially reduced. The p75 receptor displays strong affinity for the unprocessed form of NGF (pro-NGF) (Conroy and Coulson, 2022) however pro-NGF application did not affect Stmn2 protein levels (Figure 4C). We used hNGF as an internal control for human pro-NGF in this experiment. Collectively, these observations identify TrkA as the likely receptor employed by NGF to reduce Stmn2 protein.

**FIGURE 4.**
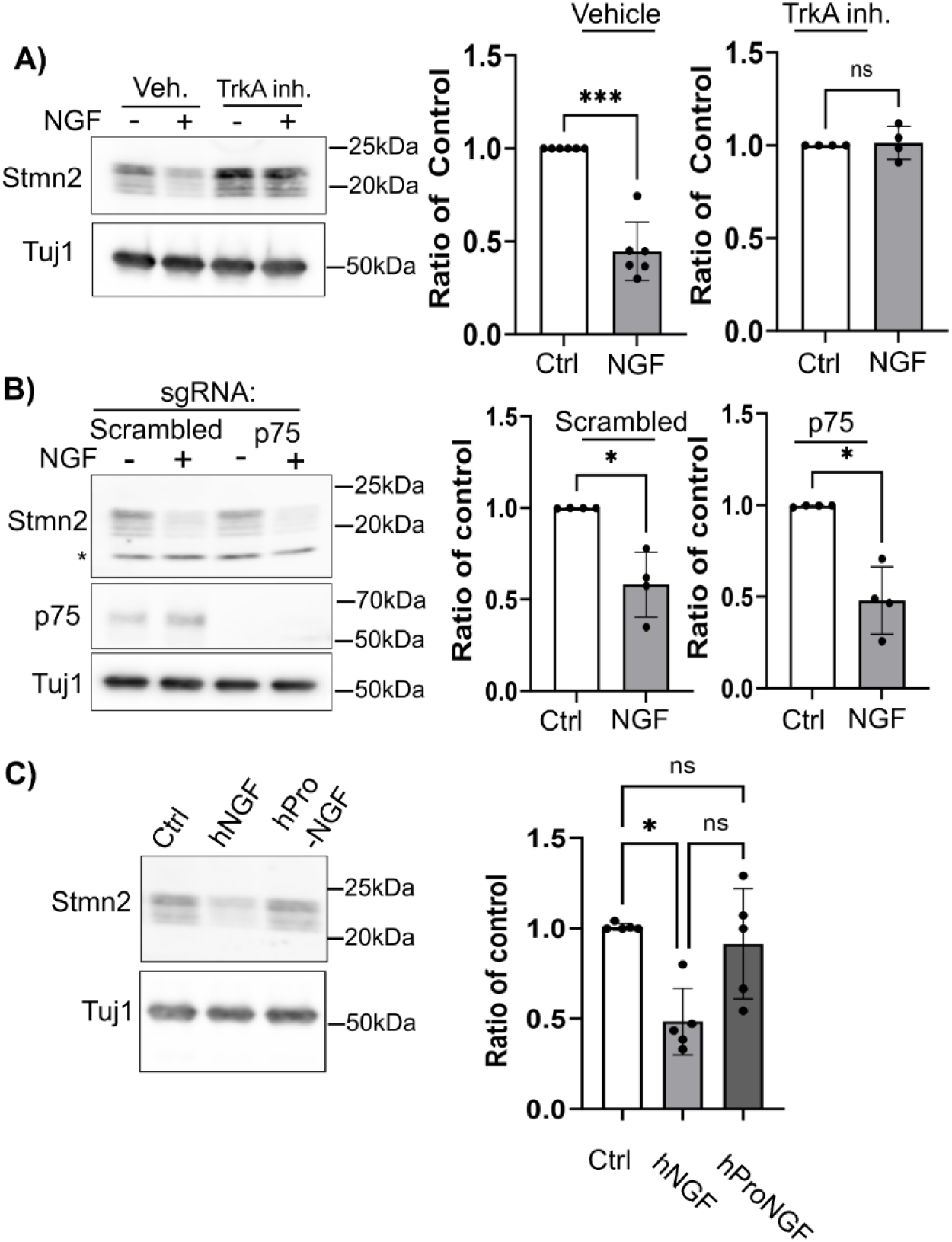
Signaling through TrkA is responsible for NGF-induced Stmn2 reduction. (A) Pretreating NGF-deprived neurons with TrkA inhibitor GW441756 (10µM) suppressed Stmn2 loss in axon-only extracts after NGF application. Quantification is shown on the right (N=4). **(B)** Neurons expressing Cas9 were transduced with lentiviruses containing scrambled sgRNA sequence or sgRNA targeting mouse p75. Western blot analysis of axon-only extracts confirms loss of p75 protein yet NGF treatment still reduced Stmn2 protein. Asterisk in Stmn2 western blot refers to non-specific band. Quantification is shown on the right (N=4). **(C)** Applying 50ng/mL human pro-NGF (hPro-NGF) did not reduce Stmn2 protein levels from axon-only extracts. Quantification is shown on the right (N=5). All statistical comparisons performed with Welch’s t-test where *p<0.05 and ***p<0.05. Error bars represent +/-1 STD.

### Analysis of signal transduction pathways downstream of TrkA

Signal transduction pathways downstream NGF-TrkA are well-established (Figure 5A). We used the NGF addback paradigm described above and manipulated each pathway with validated pharmacological inhibitors to determine which signal transduction cascade reduces axonal Stmn2 protein. Survival signaling through PI3K-Akt is particularly well-studied (Yao and Cooper, 1995; Dudek *et al*., 1997). Pharmacological inhibitors targeting PI3K and Akt (LY294002 - 20µM and Akt Inhibitor VIII - 10µM) were applied thirty minutes prior to NGF application. Axon-only extracts were collected two hours post NGF treatment. Neither inhibitor suppressed NGF-induced Stmn2 reduction (Figure 5B). Phosphorylated Akt (Ser473) was used as internal control to confirm inhibition of this pathway. Basal levels of phosphorylated Akt were undetectable in cultures undergoing chronic NGF deprivation. NGF treatment increased Akt phosphorylation and both inhibitors reduced this post-translational modification to undetectable levels indicating successful inhibition.

**FIGURE 5.**
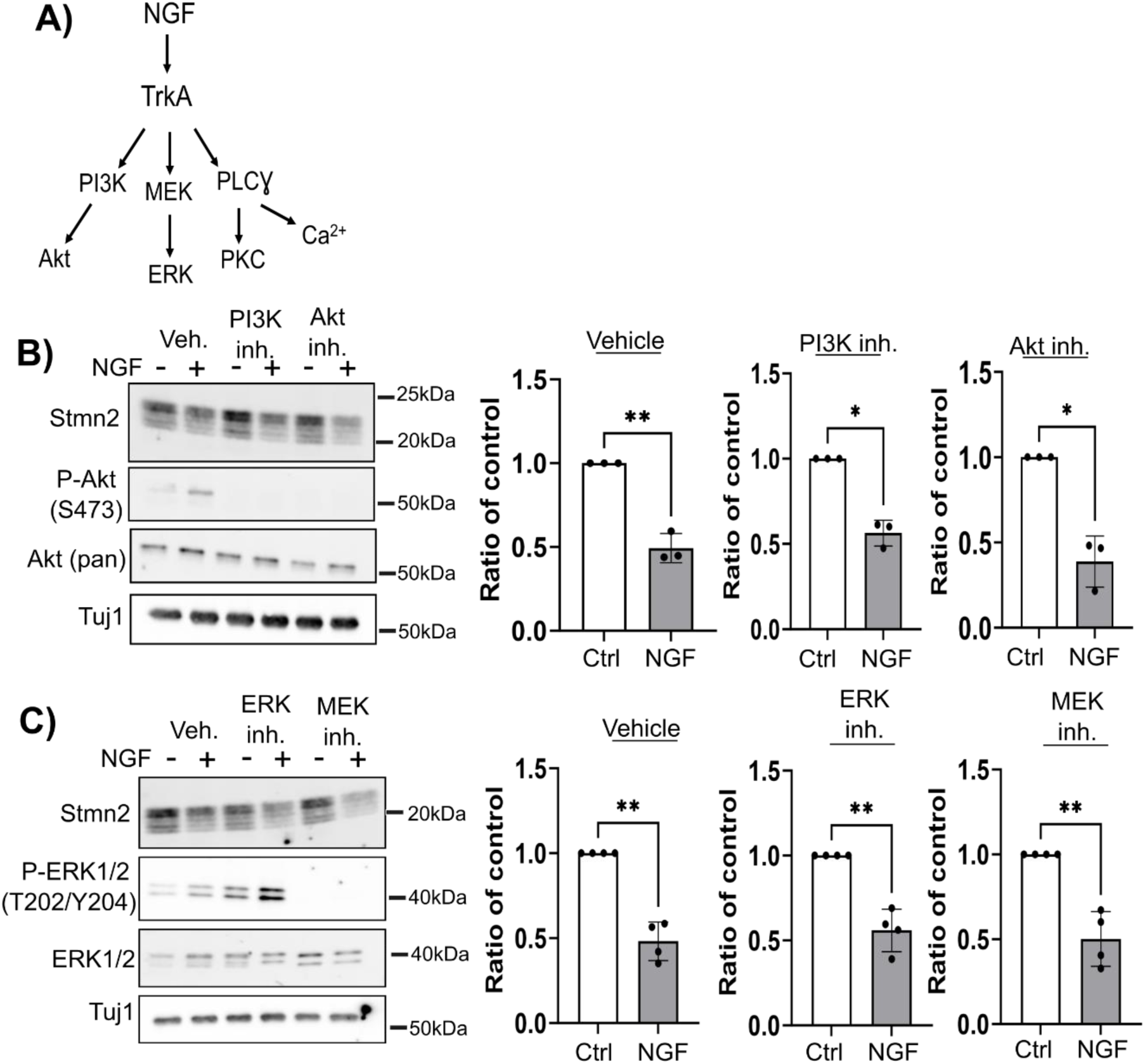
PI3K and ERK are not required for NGF-stimulated Stmn2 loss. **(A)** Canonical signaling pathways activated downstream of TrkA stimulation. **(B)** Small molecule inhibitors targeting PI3K or AKT (20µM LY294002 or 10µM AKT inhibitor VIII) did not suppress NGF-induced Stmn2 loss from axon-only extracts. Quantification is shown on the right (N=3). **(C)** Small molecule inhibitors targeting ERK1/2 or MEK1/2 (10µM Temuterkib or 10µM Selumetinib) did not suppress NGF-induced Stmn2 loss from axon-only extracts. Quantification is shown on the right (N=4). All statistical comparisons were performed with Welch’s t-test where *p<0.05 and **p<0.01. Error bars represent +/-1 STD.

The MEK/ERK pathway is a MAPK cascade activated downstream of TrkA (Thomas *et al*., 1992; Wood *et al*., 1992). Inhibitors targeting MEK1/2 or ERK1/2 (Selumetinib-10µM or Temuterkib-10µM) did not suppress NGF-induced reduction in Stmn2 protein (Figure 5C). MEK1/2 inhibition abolished ERK1/2 phosphorylation in the presence or absence of NGF. The ERK inhibitor Temuterkib elevated baseline ERK1/2 (Thr202/Tyr204) phosphorylation as well as phosphorylation provoked by NGF application. Inhibiting ERK1/2 likely suppresses activation of phosphatases responsible for turning off ERK1/2 in a negative feedback loop (Kidger and Keyse, 2016) and would account for this increase.

We next targeted phospholipase C activity (Obermeier *et al*., 1994; Stephens *et al*., 1994) with a small molecule inhibitor (U-73122) or inactive analog (U-73342) as a negative control. Phospholipase C inhibition suppressed NGF-induced Stmn2 loss while the analog displayed no effect (Figure 6A). Phospholipase C activity generates two second messengers, diacylglycerol (DAG) and inositol triphosphate (IP_3_) which stimulate PKC and Ca^2+^ influx respectively. Two broad spectrum inhibitors targeting all PKC isoforms (Go6983-10µM and sotrastaurin-10 µM) did not suppress NGF-induced Stmn2 loss (Figure 6B&C). IP_3_ stimulates opening of calcium channels at the endoplasmic reticulum. In addition to chelating intracellular Ca^2+^ with 5µM BAPTA-AM we also applied 2.5mM EGTA to chelate extracellular Ca^2+^ and account for established connections between activated TrkA and Ca^2+^ channels at the plasma membrane (Barker *et al*., 2020). These Ca^2+^ chelators were added individually and in combination thirty minutes prior to NGF application. NGF application significantly reduced Stmn2 protein under all conditions though combined EGTA/BAPTA-AM treatment displayed slight suppression (Figure 6D). Phospholipase C signaling is the leading candidate responsible for reducing Stmn2 levels in response to NGF stimulation however the mechanism is unclear.

**FIGURE 6.**
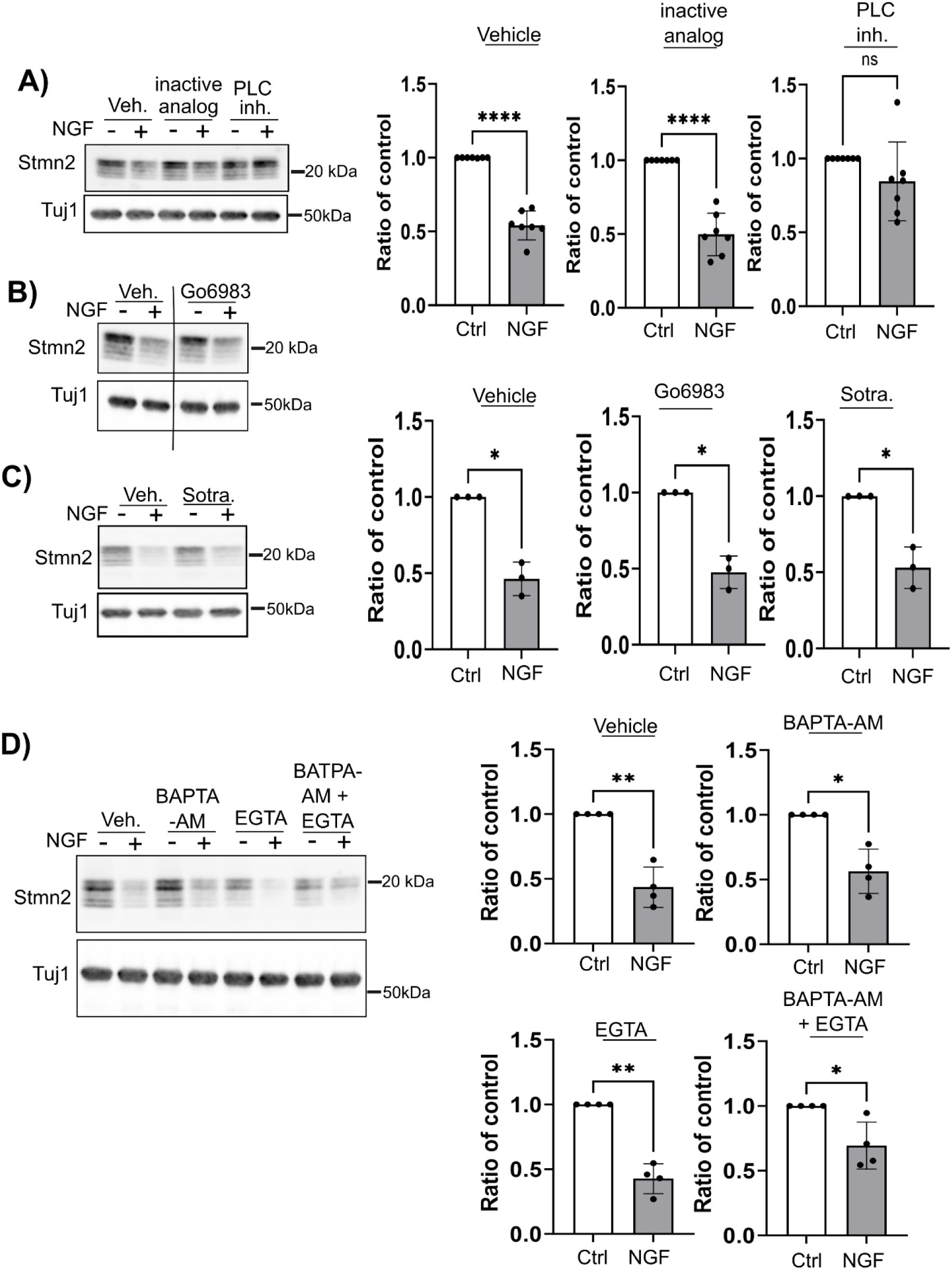
Inhibiting phospholipase C suppresses NGF-induced Stmn2 loss. **(A)** Pre-treating NGF-deprived neurons with phospholipase C inhibitor (5µM U-73122) blocked NGF-induced Stmn2 protein loss from axon-only extracts while an inactive analog (5µM U-73342) had no effect. Quantification is shown on the right (N=7). PKC inhibitors **(B)** Go6983 (10µM) and **(C)** Sotrastaurin (10µM) did not block NGF-induced Stmn2 loss from axon-only extracts. Lanes in representative western blot from Go6983 experiment were from the exposure. The full western blot is available in Supplementary Figure 2. Quantification for each treatment is shown on the right (N=3). **(D)** Calcium chelators BAPTA-AM (5µM) and EGTA (2.5mM) did not block NGF-induced Stmn2 loss from axon-only extracts. Quantification is shown on the right (N=4). All statistical comparisons were performed with Welch’s t-test where *p<0.05, **p<0.01, and ****p<0.001. Error bars represent +/-1 STD.

### NGF stimulation targets palmitoylated Stmn2 for degradation

To gain additional mechanistic insight, we next evaluated whether NGF stimulation affects axonal levels of other Stathmin proteins. Stmn1, Stmn2, and Stmn3 are phosphorylated at serine residues within a proline-rich domain (PRD) while Stmn2 and Stmn3 are also palmitoylated at an N-terminal membrane targeting domain (Chauvin and Sobel, 2015). We performed NGF addback experiments as described above and evaluated stathmin protein levels in axon-only extracts two hours after NGF stimulation. Stmn1 protein levels did not change in response to NGF treatment while Stmn3 protein levels decreased 43% (Figure 7A). To determine whether post-translational modifications are necessary for NGF-induced reduction, we expressed Venus-tagged Stmn2 variants possessing amino acid substitutions preventing either phosophorylation (Stmn2AA) or palmitolylation (Stmn2CS), chronically deprived neurons of NGF for 24 hours, then acutely stimulated neurons with NGF for two hours. Wildtype Stmn2-Venus levels decreased 40% in response to NGF (Figure 7B) while Stmn2AA and Stmn2CS levels were unaffected by NGF application.

**FIGURE 7.**
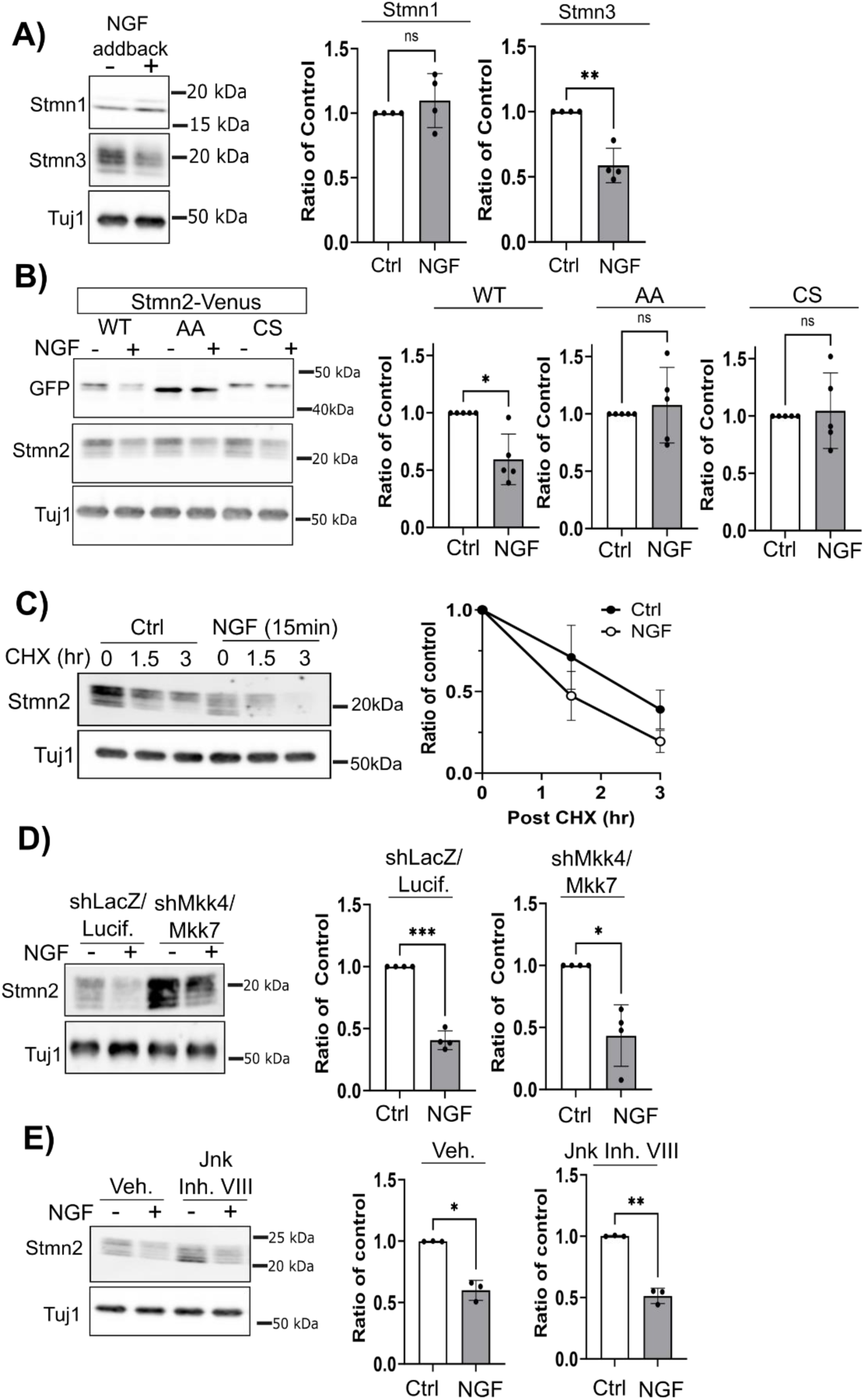
NGF stimulation triggers accelerated degradation of palmitoylated Stmn2. **(A)** NGF treatment reduces Stmn3 levels in axon-only extracts yet does not affect Stmn1 levels. Quantification is shown below (N=4). **(B)** Protein levels of Stmn2-Venus variants were measured from axon-only extracts after NGF stimulation for two hours. Amino acid substitutions in residues required for Stmn2 phosphorylation or palmitoylation block NGF-induced protein loss. Western blots of endogenous Stmn2 confirm NGF stimulation. Quantification is shown on the right (N=5). **(C)** NGF-deprived neurons were stimulated with or without NGF for fifteen minutes then neurons washed with media lacking NGF. Two hours later neurons were treated with cycloheximide (CHX - 25µg/mL) for 1.5 hour and 3 hours. Stmn2 protein levels decreased faster in axon-only extracts from NGF-stimulated neurons compared to control. Quantification is shown on the right (N=4) **(D)** Neurons were transduced with control shRNA constructs (shLacZ and shLuciferase) or shRNAs targeting Mkk4 and Mkk7. NGF application reduced Stmn2 protein from axon-only extracts under both conditions. Quantification is shown on the right (N=4). **(E)** Pretreatment with 10µM JNK inhibitor VIII did not suppress NGF-induced reduction in Stmn2 protein from axon-only extracts. Quantification is shown on the right (N=3). All statistical comparisons were performed with Welch’s T-test where *p<0.05, **p<0.01, and ***p<0.05. Error bars represent +/-1 STD.

Phosphorylation and palmitoylation regulate Stmn2 degradation (Shin *et al*., 2012; Summers *et al*., 2018). Since both post-translational modifications were necessary for NGF-induced reduction we investigated whether acute NGF stimulation affects the rate of Stmn2 turnover in axons. NGF-deprived sensory neurons were exposed to NGF for fifteen minutes to stimulate TrkA signaling. NGF washed out with fresh media lacking NGF, and neurons treated two hours later with the protein synthesis inhibitor cycloheximide (CHX). NGF pretreatment reduced Stmn2 protein levels prior to CHX application so samples were quantified as a ratio of baseline levels. In control neurons Stmn2 protein levels were reduced 20% at 1.5hr and 60% at 3hr post-CHX treatment. In contrast, NGF-pretreatment reduced Stmn2 protein levels by 60% at 1.5hr and 80% at 3hr after CHX treatment (Figure 7C).

JNK signaling promotes degradation of Stmn2 protein (Shin *et al*., 2012). We tested whether this MAPK pathway is required for NGF-stimulated degradation of Stmn2 by two methods, knocking down the upstream MAP2Ks, MKK4 and MKK7, or pretreating cells with a small molecule inhibitor to all JNK isoforms (JNK inhibitor VIII). Both manipulations elevated baseline Stmn2 protein levels as previously demonstrated (Shin *et al*., 2012; Walker *et al*., 2017) yet did not suppress NGF-induced reduction in Stmn2 (Figure 7D & E).

### Acute NGF stimulation accelerates Wallerian Degeneration

The degradation of short-lived axon maintenance factors is balanced by delivery through anterograde transport. If NGF increases Stmn2 turnover rate then depriving an axon of newly synthesized protein through axotomy should result in accelerated protein loss as well as accelerated fragmentation. To test these predictions we first performed axotomy in NGF-deprived neurons then immediately applied NGF to exclude the possibility of active transport in or out of the axon segment (Figure 8A). In NGF-deprived neurons Nmnat2 protein levels were reduced 50% one hour and 80% two hours post axotomy. Stmn2 protein reduction occurred moderately slower with levels decreasing by 40% one hour and 70% two hours post axotomy, consistent with the slightly longer half-life of this protein compared to Nmnat2. NGF application accelerated loss of both Nmnat2 and Stmn2 protein from severed axons. Nmnat2 protein levels were reduced 75% one hour and 85% two hours post axotomy while Stmn2 levels were reduced 60% one hour and 80% two hours post axotomy.

**FIGURE 8.**
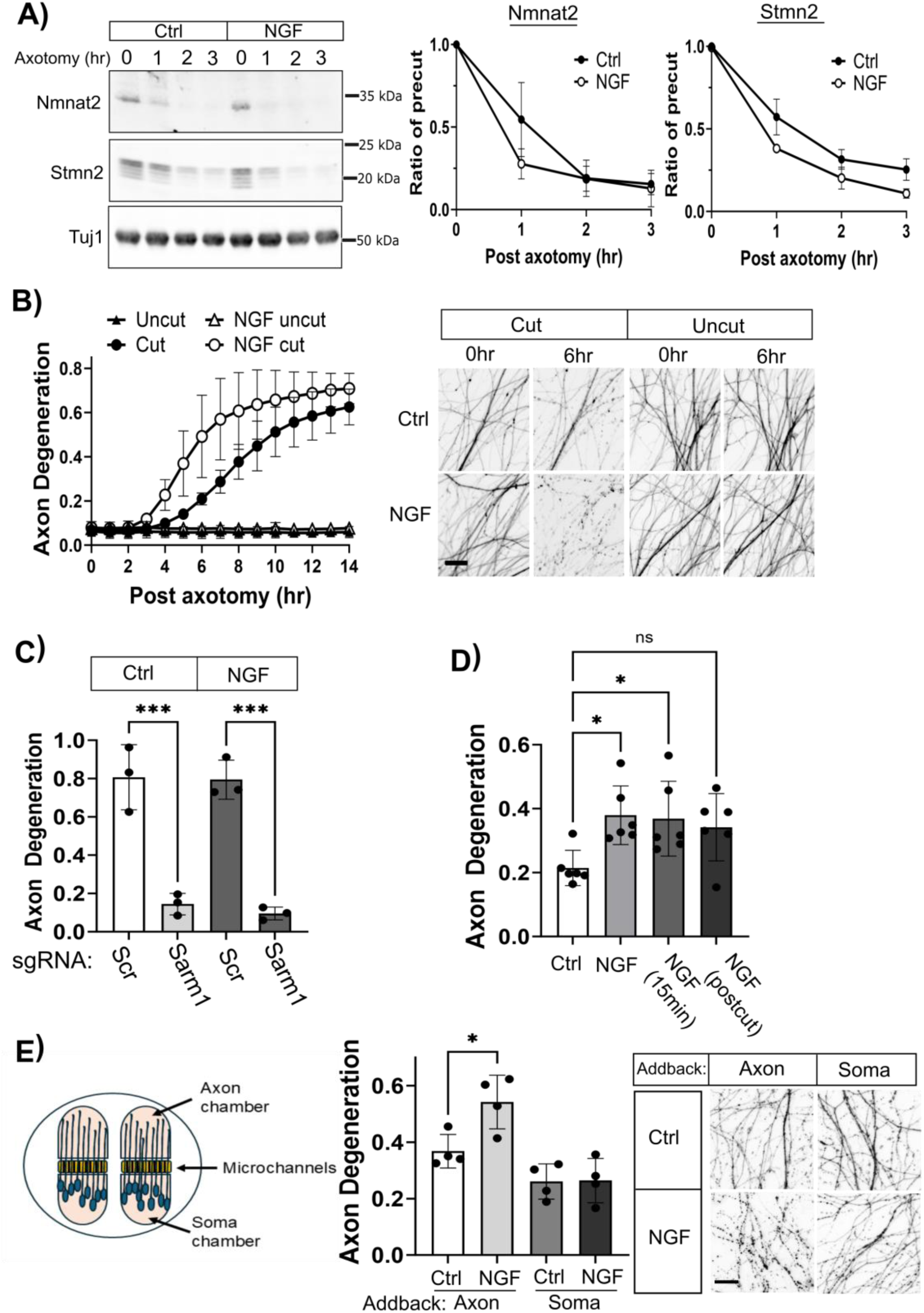
NGF stimulation accelerates Wallerian Degeneration. **(A)** Axonal Nmnat2 and Stmn2 levels decrease faster in neurons pre-treated with NGF. Quantification is shown on the right (N=4). **(B)** NGF treatment accelerated fragmentation of severed axons (N=4) with representative images from the same axon field prior to axotomy (0hr) or post axotomy (6hr) as well as axons uncut during the experimental period. **(C)** Axon degeneration measured 24hr post axotomy in Cas9-expressing neurons transduced with lentivirus expressing a scrambled (Scr) sgRNA or sgRNA targeting Sarm1, with or without NGF addback (Statistical comparisons performed with an unpaired t-test; N=3). **(D)** NGF was added to neurons at the indicated intervals. Axon degeneration was measured 10 hours after axotomy with a razor blade (Statistical comparisons performed with one-way ANOVA and post-hoc unpaired t-test; N=5). **(E)** Neurons were seeded in microfluidic devices and NGF added to either the axon chamber or the soma chamber prior to axotomy. Axon degeneration was measured 6 hours post axotomy (Statistical comparisons performed with one-way ANOVA and post-hoc unpaired t-test; N=4). Representative images of distal axons are shown on the right. For all statistical tests *p<0.05 and ***p<0.05. Error bars represent +/-1 STD. Scale bars = 20µm.

We next examined whether acute NGF stimulation accelerates fragmentation of severed axons. We chronically deprived NGF from DRGs for 24hr, applied NGF thirty minutes prior to axotomy, then measured axon degeneration over a twelve-hour period in severed axons. Blebbing and slight fragmentation were detected six hours post axotomy in control axons while NGF stimulation resulted in widespread fragmentation at this timepoint (Figure 8B). NGF treatment did not provoke axon degeneration in uncut axons as expected. CRISPR-inactivating SARM1 suppressed axon degeneration in the presence or absence of NGF, confirming NGF-accelerated fragmentation occurs through this executioner of Wallerian Degeneration (Figure 8C).

Fifteen-minute NGF pre-exposure accelerated Wallerian Degeneration to a similar extent as two-hour pre-treatment (Figure 8D), consistent with our findings that brief exposure is sufficient to reduce Nmnat2/Stmn2 protein levels. Applying NGF immediately following axotomy trended toward accelerated degeneration however did not reach statistical significance (Figure 8D). We employed microfluidic devices to ascertain whether local NGF signaling in the axon segment is sufficient to accelerate Wallerian degeneration (Figure 8E). In this experiment, NGF-deprived neurons were treated with NGF in either the axon chamber or the soma chamber and axons severed with a razor blade. We visualized severed axons six hours post axotomy when partial fragmentation is apparent in controls yet still incomplete. Applying NGF to the axon chamber enhanced axon fragmentation while treatment in the soma compartment displayed no change compared to controls (Figure 8E). Therefore, acute NGF stimulation in the axon compartment accelerates loss of maintenance factors and accelerates SARM1-dependent degeneration.

## DISCUSSION

NGF signaling promotes axonal outgrowth and sustains neuron survival. Circulating NGF increases during inflammation and is locally produced by mast cells, keratinocytes, and fibroblasts to promote wound repair in damaged tissue (Sofroniew *et al*., 2001; Minnone *et al*., 2017; Liu *et al*., 2021). Secretion and processing of NGF is balanced by degradation through extracellular proteases (Bruno and Cuello, 2006). Accordingly, axon projections experience waves of NGF exposure yet studying acute NGF stimulation in established axons from primary sensory neurons is complicated by their dependence on NGF to sustain survival. We circumvent this requirement and identify surprising consequences for NGF stimulation on proteostasis of axon maintenance factors Nmnat2 and Stmn2 as well as kinetics of Wallerian degeneration. The implications of altering both proteins during either NGF deprivation or stimulation is described below.

Local NGF deprivation provokes selective pruning of excess axonal branches without inducing neuronal cell death or degeneration of the primary axonal projection (Geden *et al*., 2019). Nmnat enzymes display antagonistic roles on axon regeneration (Chen *et al*., 2016; Kim *et al*., 2018) and increasing Nmnat2 local NGF deprivation is consistent with a role in suppressing axonal outgrowth. Boosting Nmnat2 would also restrain Sarm1 activation and prevent widespread dismantling of the primary axon projection or destructive signaling to the immune system (Gilley and Coleman, 2010; Gilley *et al*., 2015; Hsu *et al*., 2021; Dingwall *et al*., 2022). Nmnat2 is the terminal enzyme in a NAD^+^ salvage pathway and augmenting local Nmnat2 could alter activity of NAD^+^-dependent enzymes like SIRTs which regulate microtubule dynamics (Harkcom *et al*., 2014). Microtubule destabilizing factors such Kif2A are critical for disassembling microtubule populations in NGF-deprived axons and remodeling skin innervation *in vivo* (Maor-Nof *et al*., 2013; Dey *et al*., 2023). Increasing Stmn2 protein would sequester heterotubulin dimers thereby reducing the pool available for microtubule polymerization (Chauvin and Sobel, 2015), likewise consistent with suppressing axonal outgrowth in branches undergoing pruning. Further studies will need to determine whether fluctuations in Nmnat2 abundance elicit corresponding changes in local NAD^+^ generation and whether NAD^+^ hydrolysis through Sarm1 is connected to axonal remodeling.

NGF-TrkA activation promote axonal outgrowth and collateral branch formation in part through actin polymerization (Spillane *et al*., 2012) and local debundling of microtubules at branch points (Ketschek *et al*., 2015). Microtubules infiltrate a subpopulation of immature of collateral branches supporting physical stability and maturation through motor-driven delivery of vesicles and mitochondria (Armijo-Weingart and Gallo, 2017). NGF signaling reduces Stmn2 and Stmn3 abundance and would facilitate microtubule polymerization into collateral branches. NGF stimulation did not affect axonal Stmn1 protein levels yet NGF does stimulate Stmn1 phosphorylation which would inhibit Stathmin:tubulin interaction (Doye *et al*., 1990). Mutagenesis studies indicate both palmitoylation and phosphorylation are necessary for NGF-induced Stmn2 loss. Palmitoylated Stmn2 regulates membrane trafficking through unclear mechanisms (Mahapatra *et al*., 2008; Wang *et al*., 2013), raising the possibility that Stmn2 subpopulations control vesicle exocytosis at axonal branch points. Nmnat2 might be a bystander in NGF-induced reduction of Stmn2 as these axon maintenance factors co-localize on vesicles and undergo degradation through some parallel mechanisms (Summers *et al*., 2018). Alternatively, Nmnats regulate synaptic activity (Zang *et al*., 2013; Russo *et al*., 2019) and presynaptic remodeling might depend on modifying NAD^+^ homeostasis or reducing Nmnat chaperone activity.

Mature NGF promotes survival through preferential binding to the high affinity receptor TrkA while proNGF signaling through the lower affinity receptor p75 increases neurodegeneration (Mufson *et al*., 2019). MAPK signaling through JNK is a known effector of p75 however CRISPR-editing and pharmacology strongly indicate these pathways are not involved and TrkA is the relevant receptor. Our pharmacological studies point to phospholipase C as the effector for TrkA-dependent Stmn2 loss however we could not pinpoint which second messenger generated by phospholipase C (DAG or IP_3_) is responsible. Blocking Ca^2+^ influx through intracellular and extracellular sources showed promise however did not convincingly suppress NGF-induced Stmn2 reduction. DAG can be hydrolyzed into other metabolites with signaling functions beyond PKC activation (Eichmann and Lass, 2015). Connections between phospholipid metabolism and degradation of palmitoylated Stmn2 warrant future investigation.

Intersections between developmental axon pruning and neurodegeneration have intrigued scientists for many decades (Raff *et al*., 2002; Yaron and Schuldiner, 2016; Geden *et al*., 2019). Death Receptor 6 promotes axon degeneration in response to both NGF deprivation and axotomy (Gamage *et al*., 2017). Wnk kinases regulate axon branching during development as well as axon maintenance in adulthood through additive Sarm1 suppression with Nmnat enzymes (Izadifar *et al*., 2021). Even though our observations suggest NGF signaling antagonizes axonal maintenance proteins, the therapeutic potential of local NGF application is well-supported in numerous preclinical disease models across multiple decades (Mobley, 1989; Lambiase *et al*., 2009; Amadoro *et al*., 2021). Rather, our study suggests NGF signaling primes axon compartments toward regrowth and repair at the expense of transient susceptibility to Sarm1-dependent degeneration. NGF biology continues to offer many surprises with more discoveries waiting in the future.

## Methods

### Plasmids and reagents

Bcl-Xl and Stmn2-Venus expression constructs were described previously (Thornburg-Suresh *et al*., 2023). Myristoylated Scarlet (myrScarlet) was generated by PCR amplification from a plasmid backbone containing the Scarlet open reading frame (a gift from Erik Dent, Addgene plasmid#125138; http://n2t.net/addgene:125138; RRID:Addgene_125138) and Gibson cloning with a 5’ insertion encoding an eight amino acid myristoylation sequence derived from human Src into a lentiviral expression backbone with the human ubiquitin promoter. Myristoylated Akt1 was a gift from Heng Zhao (Addgene #53583; http://n2t.net/addgene:53583; RRID:Addgene_53583).’ In CRISPR-editing studies two independent sgRNAs targeting mouse p75 (NGFR) or Sarm1 were designed with CRISPick (Broad Institute) and ligated into BsmBI-digested Lentiguide plasmid backbone. Sequences for p75 targeting sgRNAs were #1 5’ ACAGGCATGTACACCCACA 3’ and sgRNA #2 5’GAGTATGTCCGCTCCCTGT 3’. Sequence for Sarm1-targeting sgRNA was 5’ TCGCGAAGTGTCGCCCGGAG 3’. Two scramble sgRNAs were used as controls, #1 5’ CGTCGCCGGCGAATTGACGG 3’ and #2 5’ CGCGGCAGCCGGTAGCTATG 3’. Knockdown constructs (shLacZ, shLuciferase, shMkk4 and shMkk7) are previously published (Walker *et al*., 2017). Media components and their sources are listed here. DRG sensory neurons were cultured in phenol-red free Neurobasal media (Gibco) supplemented with 2mM glutamine, 10 U/mL penicillin/streptomycin, 2% B27 supplement (all from Gibco), 50ng/mL mouse 2.5S NGF (Alomone Labs), and 1mM 5-fluorodeoxyuridine/1mM uridine (Thermofisher). HEK cells were cultured in DMEM (4.5g/L glucose; Corning) supplemented with heat-inactivated Fetal Bovine Serum (Corning), 2mM L-glutamine, and penicillin/streptomycin (10U/mL). Recombinant human beta NGF, proNGF, and BDNF were from Alomone labs. Chemicals utilized in this study and their source are listed here: Sotrastaurin (Medchemexpress), BAPTA-AM (Biotium), EGTA Research Products International), cycloheximide (Thermo Scientfic) and the following were from Cayman Chemical, JNK Inhibitor VIII, AKT inhibitor VIII, Go 6983, Selumitinib, Temuterkib, LY294002, U-73122, U-73342. Fresh aliquots were used for each experimental replicate.

### Culture of primary embryonic sensory neurons and lentiviral transduction

Pregnant CD1 mice were from Charles River Laboratory. Dorsal root ganglia (DRGs) were dissected from E13.5 embryos (a mixture of both male and female) and spotted on plates precoated with poly-d-lysine (Sigma) and laminin (Gibco). Neurons were cultured in neurobasal media prepared as described above containing NGF for six days until NGF-manipulating experiments were initiated. Lentivirus was prepared as previously described (Gerdts *et al*., 2011). Briefly, HEK293 cells were co-transfected with vesicular stomatitis glycoprotein, the lentiviral packaging plasmid PspAX2, and an expression plasmid under control of the human ubiquitin promoter. Media containing lentivirus was collected two days later, dead cells removed by centrifugation, and supernatant stored in aliquots at −80°C. Lentivirus expressing Bcl-xL was applied to sensory neurons on Day *in vitro* 2 (DIV2) while lentivirus expressing Stmn2-Venus constructs was applied on DIV5. For axon degeneration and microscopy studies, DRG sensory neurons were transduced on DIV2 with myristoylated-Scarlet to label axons. In CRISPR-editing experiments lentivirus expressing Cas9 and sgRNAs were added on DIV1. Experiments were performed on DIV7 and DIV8.

### NGF deprivation

DRG sensory neurons underwent three media changes with neurobasal media containing all the components listed above except NGF. In the final media change neurons were supplied with media +/- NGF (50ng/mL). For studies of acute NGF deprivation described in Figure 1 and Figure 2, NGF-lacking media also contained anti-sera to NGF (Sigma, 1:5000, RRID:AB_477660). This antisera was omitted in experiments evaluating NGF addback described in Figure 1F and Figures 3 – 8. For these experiments DRG neurons were washed three times with media lacking NGF twenty-four hours prior to re-applying NGF and analysis.

### Measurements of axon degeneration

For timelapse studies, DRG sensory neurons were spotted in 96-well dishes and transduced with myristoylated-Scarlet to label neuronal membranes. Axons were severed with a razor blade under the indicated experimental conditions and distal axon segments visualized once an hour with an automated microscope (either a Cytation 5 or Lionheart Imager from Agilent). Axon degeneration score was calculated from each image using an ImageJ macro that measures fragmented axon area from a field based on a pre-determined circulatory score assigned to each object (Gerdts *et al*., 2011). In studies with microfluidic devices DRG sensory neurons were fixed in 3.7% formaldehyde six-hours post axotomy and images collected manually with a Lecia DM IL inverted microscope. Quantification of axon degeneration from these images was performed with the same ImageJ macro described above.

### Immunofluorescence detection of endogenous Stmn2

DRG sensory neurons were seeded in 35mm dishes (World Precision Instruments) or microfluidic devices (eNuvio). Cells were fixed in 3.7% formaldehyde and subsequently blocked/permeabilized in phosphate buffered saline (PBS) with 0.05% triton-x and 2.5% goat serum for 15 minutes at room temperature. Specimens were incubated overnight with anti-Stmn2 antibody (Proteintech, 1:250, RRID:AB_2197283) prepared in blocking buffer, washed three times in PBS, and incubated for one hour with secondary antibody (Alexa488-conjugated anti-Rabbit, 1:500). Following three washes in PBS, specimens, Stmn2 and myristoylated-Scarlet were visualized with an Echo spinning disk confocal microscope. Z-stacks were collected for each field. Z-projections based on max intensity were used in quantification. Myristoylated-Scarlet images were used to generate a mask for quantifying mean fluorescence intensity from corresponding Stmn2 immunofluorescence images. At least six distal axon fields were collected from each experimental replicate derived from independent mouse litters.

### Protein analysis from axon-only extracts

DRG sensory neurons were seeded in concentrated spot cultures within 12-well dishes. At the time of protein extraction, cells were washed in cold PBS and a razor blade was used to cut around the soma so a pipet tip could dislodge the soma cluster. The remaining axon field was lysed in cold RIPA buffer (50mM Tris-HCl pH 7.4, 150mM NaCl, 1mM EDTA, 1% Triton X-100, 0.5% sodium deoxycholate, and 0.1% sodium dodecyl sulfate) supplemented with fresh protease inhibitor and phosphatase inhibitor (Halt 100x cocktail, Thermo Scientific). Extracts were pre-cleared of cell debris by centrifugation (5,000xg for 5min). Supernatants were transferred into sample buffer (65.2mM Tris-HCl pH 6.8, 2% SDS, 10% glycerol, 8% beta-mercaptoethanol, 0.025% bromophenol blue with fresh beta-mercaptoethanol. Samples were boiled five minutes and separated by SDS-PAGE followed by western immunoblotting. The following antibodies were used for western immunoblotting: Stmn2 (Proteintech, RRID:AB_2197283, 1:1000), Stmn1 (Cell Signaling; RRID:AB_2798284; 1:1,000), Stmn3 (Proteintech; RRID:AB_2197399; 1:1,000), anti-GFP (Thermo Fisher; RRID:AB_221569; 1:1,000), ERK1/2 (Cell Signaling; RRID:AB_390779, 1:1000), phosphoERK1/2 Thr220/Tyr204 (Cell Signaling; RRID:AB_2315112, 1:1000), Akt (pan) (Cell Signaling; RRID:915783, 1:1000), phosphoAkt Ser473 (Cell Signaling; RRID:2315049, 1:000). Primary antibodies were detected with dye-conjugated secondary antibodies (Li-Cor anti-mouse 800CW RRID:AB_2687825 and Thermo Scientific anti-Rabbit Alexa Fluor 680 RRID:AB2536103, 1:5000) and visualized with a Li-Cor® Odyssey Fc Imaging system.

## Acknowledgements

Research described in this manuscript was supported by funds from the National Institutes of Health to D.W.S. (RO1NS126191). We appreciate thoughtful comments from members of the Summers during preparation of this manuscript.

## Author Contributions

J.A.D. participated in experimental design, data collection, data analysis, and writing of this manuscript. M.A. and L.M. participated in data collection. D.W.S. participated in experimental design, data collection, data analysis, and writing of this manuscript.

---

**Supplementary Figure 1.**
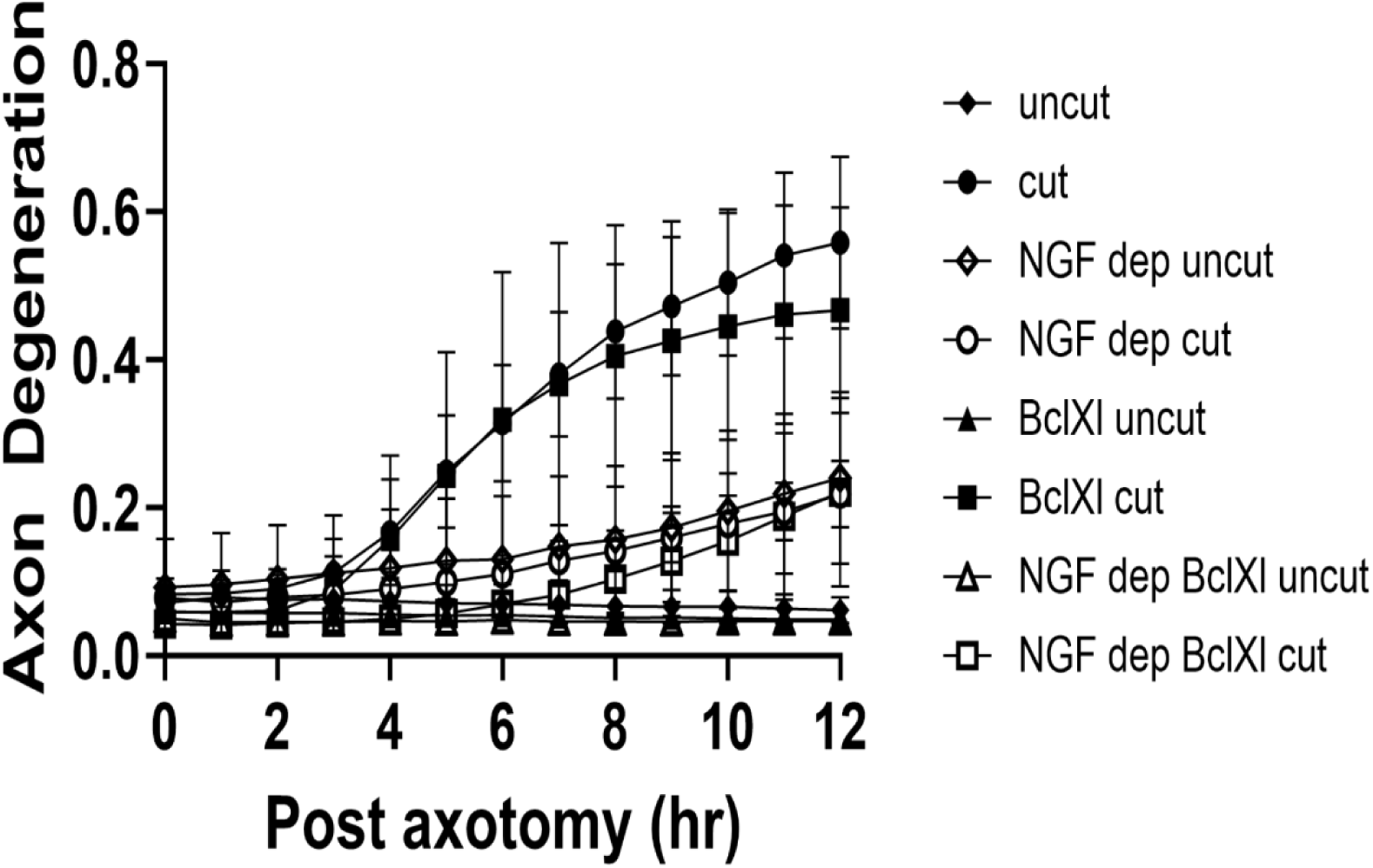
Bcl-xL overexpression does not affect axon protection afforded by NGF-deprivation. DRG sensory neurons transduced with lentivirus containing an empty vector or Bcl-xL expression construct. NGF-deprivation was performed as described in the main text and axons severed with a razor blade. Uncut axons were used as controls. Degeneration of distal axons was quantified over time (N=4). Error bars represent +/- 1 STD.

**Supplementary Figure 2.**
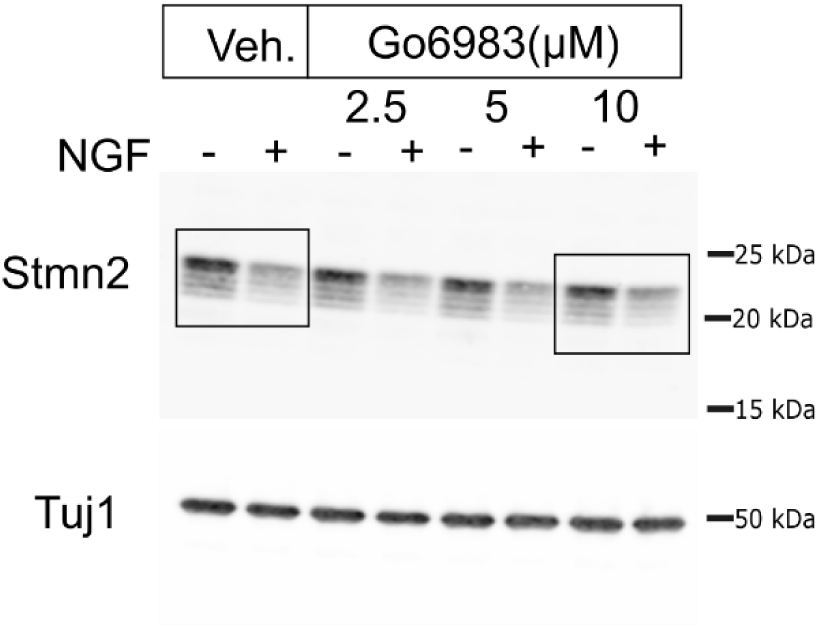
The PKC inhibitor Go6983 does not suppress NGF-induced Stmn2 reduction. Western blot from Figure 6B with cropped lanes outlined.

## References

Amadoro, G., Latina, V., Balzamino, B.O., Squitti, R., Varano, M., Calissano, P., and Micera, A. (2021). Nerve Growth Factor-Based Therapy in Alzheimer’s Disease and Age-Related Macular Degeneration. Front Neurosci 15, 735928.

Armijo-Weingart, L., and Gallo, G. (2017). It takes a village to raise a branch: Cellular mechanisms of the initiation of axon collateral branches. Mol Cell Neurosci 84, 36–47.

Arthur-Farraj, P., and Coleman, M.P. (2021). Lessons from Injury: How Nerve Injury Studies Reveal Basic Biological Mechanisms and Therapeutic Opportunities for Peripheral Nerve Diseases. Neurotherapeutics 18, 2200–2221.

Barker, P.A., Mantyh, P., Arendt-Nielsen, L., Viktrup, L., and Tive, L. (2020). Nerve Growth Factor Signaling and Its Contribution to Pain. J Pain Res 13, 1223–1241.

Bruno, M.A., and Cuello, A.C. (2006). Activity-dependent release of precursor nerve growth factor, conversion to mature nerve growth factor, and its degradation by a protease cascade. Proc Natl Acad Sci U S A 103, 6735–6740.

Chauvin, S., and Sobel, A. (2015). Neuronal stathmins: a family of phosphoproteins cooperating for neuronal development, plasticity and regeneration. Prog Neurobiol 126, 1–18.

Chen, L., Nye, D.M., Stone, M.C., Weiner, A.T., Gheres, K.W., Xiong, X., Collins, C.A., and Rolls, M.M. (2016). Mitochondria and Caspases Tune Nmnat-Mediated Stabilization to Promote Axon Regeneration. PLoS Genet 12, e1006503.

Coleman, M.P., and Hoke, A. (2020). Programmed axon degeneration: from mouse to mechanism to medicine. Nat Rev Neurosci 21, 183–196.

Conroy, J.N., and Coulson, E.J. (2022). High-affinity TrkA and p75 neurotrophin receptor complexes: A twisted affair. J Biol Chem 298, 101568.

de Leon, A., Gibon, J., and Barker, P.A. (2021). NGF-Dependent and BDNF-Dependent DRG Sensory Neurons Deploy Distinct Degenerative Signaling Mechanisms. eNeuro 8.

Deppmann, C.D., Mihalas, S., Sharma, N., Lonze, B.E., Niebur, E., and Ginty, D.D. (2008). A model for neuronal competition during development. Science 320, 369–373.

Dey, S., Barkai, O., Gokhman, I., Suissa, S., Haffner-Krausz, R., Wigoda, N., Feldmesser, E., Ben-Dor, S., Kovalenko, A., Binshtok, A., and Yaron, A. (2023). Kinesin family member 2A gates nociception. Cell Rep 42, 113257.

Dingwall, C.B., Strickland, A., Yum, S.W., Yim, A.K., Zhu, J., Wang, P.L., Yamada, Y., Schmidt, R.E., Sasaki, Y., Bloom, A.J., DiAntonio, A., and Milbrandt, J. (2022). Macrophage depletion blocks congenital SARM1-dependent neuropathy. J Clin Invest 132.

Doye, V., Boutterin, M.C., and Sobel, A. (1990). Phosphorylation of stathmin and other proteins related to nerve growth factor-induced regulation of PC12 cells. J Biol Chem 265, 11650–11655.

Dudek, H., Datta, S.R., Franke, T.F., Birnbaum, M.J., Yao, R., Cooper, G.M., Segal, R.A., Kaplan, D.R., and Greenberg, M.E. (1997). Regulation of neuronal survival by the serine-threonine protein kinase Akt. Science 275, 661–665.

Eichmann, T.O., and Lass, A. (2015). DAG tales: the multiple faces of diacylglycerol--stereochemistry, metabolism, and signaling. Cell Mol Life Sci 72, 3931–3952.

Figley, M.D., and DiAntonio, A. (2020). The SARM1 axon degeneration pathway: control of the NAD(+) metabolome regulates axon survival in health and disease. Curr Opin Neurobiol 63, 59–66.

Figley, M.D., Gu, W., Nanson, J.D., Shi, Y., Sasaki, Y., Cunnea, K., Malde, A.K., Jia, X., Luo, Z., Saikot, F.K., Mosaiab, T., Masic, V., Holt, S., Hartley-Tassell, L., McGuinness, H.Y., Manik, M.K., Bosanac, T., Landsberg, M.J., Kerry, P.S., Mobli, M., Hughes, R.O., Milbrandt, J., Kobe, B., DiAntonio, A., and Ve, T. (2021). SARM1 is a metabolic sensor activated by an increased NMN/NAD(+) ratio to trigger axon degeneration. Neuron 109, 1118–1136 e1111.

Gamage, K.K., Cheng, I., Park, R.E., Karim, M.S., Edamura, K., Hughes, C., Spano, A.J., Erisir, A., and Deppmann, C.D. (2017). Death Receptor 6 Promotes Wallerian Degeneration in Peripheral Axons. Curr Biol 27, 890–896.

Garcia, I., Martinou, I., Tsujimoto, Y., and Martinou, J.C. (1992). Prevention of programmed cell death of sympathetic neurons by the bcl-2 proto-oncogene. Science 258, 302–304.

Geden, M.J., Romero, S.E., and Deshmukh, M. (2019). Apoptosis versus axon pruning: Molecular intersection of two distinct pathways for axon degeneration. Neurosci Res 139, 3–8.

Geisler, S. (2024). Augustus Waller’s foresight realized: SARM1 in peripheral neuropathies. Curr Opin Neurobiol 87, 102884.

Gerdts, J., Sasaki, Y., Vohra, B., Marasa, J., and Milbrandt, J. (2011). Image-based screening identifies novel roles for IkappaB kinase and glycogen synthase kinase 3 in axonal degeneration. J Biol Chem 286, 28011–28018.

Gilley, J., and Coleman, M.P. (2010). Endogenous Nmnat2 Is an Essential Survival Factor for Maintenance of Healthy Axons. PLoS Biology 8, e1000300.

Gilley, J., Orsomando, G., Nascimento-Ferreira, I., and Coleman, Michael P. (2015). Absence of SARM1 Rescues Development and Survival of NMNAT2-Deficient Axons. Cell Reports 10, 1974–1981.

Harkcom, W.T., Ghosh, A.K., Sung, M.S., Matov, A., Brown, K.D., Giannakakou, P., and Jaffrey, S.R. (2014). NAD+ and SIRT3 control microtubule dynamics and reduce susceptibility to antimicrotubule agents. Proc Natl Acad Sci U S A 111, E2443–2452.

Hempstead, B.L., Martin-Zanca, D., Kaplan, D.R., Parada, L.F., and Chao, M.V. (1991). High-affinity NGF binding requires coexpression of the trk proto-oncogene and the low-affinity NGF receptor. Nature 350, 678–683.

Holland, S.M., Collura, K.M., Ketschek, A., Noma, K., Ferguson, T.A., Jin, Y., Gallo, G., and Thomas, G.M. (2016). Palmitoylation controls DLK localization, interactions and activity to ensure effective axonal injury signaling. Proceedings of the National Academy of Sciences 113, 763–768.

Hsu, J.M., Kang, Y., Corty, M.M., Mathieson, D., Peters, O.M., and Freeman, M.R. (2021). Injury-Induced Inhibition of Bystander Neurons Requires dSarm and Signaling from Glia. Neuron 109, 473–487 e475.

Izadifar, A., Courchet, J., Virga, D.M., Verreet, T., Hamilton, S., Ayaz, D., Misbaer, A., Vandenbogaerde, S., Monteiro, L., Petrovic, M., Sachse, S., Yan, B., Erfurth, M.L., Dascenco, D., Kise, Y., Yan, J., Edwards-Faret, G., Lewis, T., Polleux, F., and Schmucker, D. (2021). Axon morphogenesis and maintenance require an evolutionary conserved safeguard function of Wnk kinases antagonizing Sarm and Axed. Neuron 109, 2864–2883 e2868.

Kaplan, D.R., and Stephens, R.M. (1994). Neurotrophin signal transduction by the Trk receptor. J Neurobiol 25, 1404–1417.

Ketschek, A., Jones, S., Spillane, M., Korobova, F., Svitkina, T., and Gallo, G. (2015). Nerve growth factor promotes reorganization of the axonal microtubule array at sites of axon collateral branching. Dev Neurobiol 75, 1441–1461.

Khan, N., and Smith, M.T. (2015). Neurotrophins and Neuropathic Pain: Role in Pathobiology. Molecules 20, 10657–10688.

Kidger, A.M., and Keyse, S.M. (2016). The regulation of oncogenic Ras/ERK signalling by dual-specificity mitogen activated protein kinase phosphatases (MKPs). Semin Cell Dev Biol 50, 125–132.

Kim, K.W., Tang, N.H., Piggott, C.A., Andrusiak, M.G., Park, S., Zhu, M., Kurup, N., Cherra, S.J., 3rd, Wu, Z., Chisholm, A.D., and Jin, Y. (2018). Expanded genetic screening in Caenorhabditis elegans identifies new regulators and an inhibitory role for NAD(+) in axon regeneration. Elife 7.

Klim, J.R., Williams, L.A., Limone, F., Guerra San Juan, I., Davis-Dusenbery, B.N., Mordes, D.A., Burberry, A., Steinbaugh, M.J., Gamage, K.K., Kirchner, R., Moccia, R., Cassel, S.H., Chen, K., Wainger, B.J., Woolf, C.J., and Eggan, K. (2019). ALS-implicated protein TDP-43 sustains levels of STMN2, a mediator of motor neuron growth and repair. Nat Neurosci 22, 167–179.

Ko, K.W., Devault, L., Sasaki, Y., Milbrandt, J., and DiAntonio, A. (2021). Live imaging reveals the cellular events downstream of SARM1 activation. Elife 10.

Kohn, A.D., Takeuchi, F., and Roth, R.A. (1996). Akt, a pleckstrin homology domain containing kinase, is activated primarily by phosphorylation. J Biol Chem 271, 21920–21926.

Krauss, R., Bosanac, T., Devraj, R., Engber, T., and Hughes, R.O. (2020). Axons Matter: The Promise of Treating Neurodegenerative Disorders by Targeting SARM1-Mediated Axonal Degeneration. Trends in Pharmacological Sciences 41, 281–293.

Lambiase, A., Aloe, L., Centofanti, M., Parisi, V., Bao, S.N., Mantelli, F., Colafrancesco, V., Manni, G.L., Bucci, M.G., Bonini, S., and Levi-Montalcini, R. (2009). Experimental and clinical evidence of neuroprotection by nerve growth factor eye drops: Implications for glaucoma. Proc Natl Acad Sci U S A 106, 13469–13474.

Levi-Montalcini, R., and Booker, B. (1960). Destruction of the Sympathetic Ganglia in Mammals by an Antiserum to a Nerve-Growth Protein. Proc Natl Acad Sci U S A 46, 384–391.

Lewin, G.R., Ritter, A.M., and Mendell, L.M. (1993). Nerve growth factor-induced hyperalgesia in the neonatal and adult rat. J Neurosci 13, 2136–2148.

Liu, Z., Wu, H., and Huang, S. (2021). Role of NGF and its receptors in wound healing (Review). Exp Ther Med 21, 599.

Luo, L., and O’Leary, D.D. (2005). Axon retraction and degeneration in development and disease. Annu Rev Neurosci 28, 127–156.

Mahapatra, N.R., Taupenot, L., Courel, M., Mahata, S.K., and O’Connor, D.T. (2008). The *trans* -Golgi Proteins SCLIP and SCG10 Interact with Chromogranin A To Regulate Neuroendocrine Secretion. Biochemistry 47, 7167–7178.

Maor-Nof, M., Homma, N., Raanan, C., Nof, A., Hirokawa, N., and Yaron, A. (2013). Axonal pruning is actively regulated by the microtubule-destabilizing protein kinesin superfamily protein 2A. Cell Rep 3, 971–977.

Meeker, R.B., and Williams, K.S. (2015). The p75 neurotrophin receptor: at the crossroad of neural repair and death. Neural Regen Res 10, 721–725.

Melamed, Z., Lopez-Erauskin, J., Baughn, M.W., Zhang, O., Drenner, K., Sun, Y., Freyermuth, F., McMahon, M.A., Beccari, M.S., Artates, J.W., Ohkubo, T., Rodriguez, M., Lin, N., Wu, D., Bennett, C.F., Rigo, F., Da Cruz, S., Ravits, J., Lagier-Tourenne, C., and Cleveland, D.W. (2019). Premature polyadenylation-mediated loss of stathmin-2 is a hallmark of TDP-43-dependent neurodegeneration. Nat Neurosci 22, 180–190.

Minnone, G., De Benedetti, F., and Bracci-Laudiero, L. (2017). NGF and Its Receptors in the Regulation of Inflammatory Response. Int J Mol Sci 18.

Mobley, W.C. (1989). Nerve growth factor in Alzheimer’s disease: to treat or not to treat? Neurobiol Aging 10, 578–580; discussion 588-590.

Mufson, E.J., Counts, S.E., Ginsberg, S.D., Mahady, L., Perez, S.E., Massa, S.M., Longo, F.M., and Ikonomovic, M.D. (2019). Nerve Growth Factor Pathobiology During the Progression of Alzheimer’s Disease. Front Neurosci 13, 533.

Niu, J., Holland, S.M., Ketschek, A., Collura, K.M., Hesketh, N.L., Hayashi, T., Gallo, G., and Thomas, G.M. (2022). Palmitoylation couples the kinases DLK and JNK3 to facilitate prodegenerative axon-to-soma signaling. Science Signaling 15, eabh2674.

Obermeier, A., Bradshaw, R.A., Seedorf, K., Choidas, A., Schlessinger, J., and Ullrich, A. (1994). Neuronal differentiation signals are controlled by nerve growth factor receptor/Trk binding sites for SHC and PLC gamma. EMBO J 13, 1585–1590.

Petty, B.G., Cornblath, D.R., Adornato, B.T., Chaudhry, V., Flexner, C., Wachsman, M., Sinicropi, D., Burton, L.E., and Peroutka, S.J. (1994). The effect of systemically administered recombinant human nerve growth factor in healthy human subjects. Ann Neurol 36, 244–246.

Raff, M.C., Whitmore, A.V., and Finn, J.T. (2002). Axonal self-destruction and neurodegeneration. Science 296, 868–871.

Russo, A., Goel, P., Brace, E.J., Buser, C., Dickman, D., and DiAntonio, A. (2019). The E3 ligase Highwire promotes synaptic transmission by targeting the NAD-synthesizing enzyme dNmnat. EMBO Rep 20.

Saxena, S., and Caroni, P. (2007). Mechanisms of axon degeneration: From development to disease. Progress in Neurobiology 83, 174–191.

Sengupta Ghosh, A., Wang, B., Pozniak, C.D., Chen, M., Watts, R.J., and Lewcock, J.W. (2011). DLK induces developmental neuronal degeneration via selective regulation of proapoptotic JNK activity. Journal of Cell Biology 194, 751–764.

Shin, J.E., Miller, B.R., Babetto, E., Cho, Y., Sasaki, Y., Qayum, S., Russler, E.V., Cavalli, V., Milbrandt, J., and DiAntonio, A. (2012). SCG10 is a JNK target in the axonal degeneration pathway. Proc Natl Acad Sci U S A 109, E3696–3705.

Simon, D.J., Pitts, J., Hertz, N.T., Yang, J., Yamagishi, Y., Olsen, O., Tesic Mark, M., Molina, H., and Tessier-Lavigne, M. (2016). Axon Degeneration Gated by Retrograde Activation of Somatic Pro-apoptotic Signaling. Cell 164, 1031–1045.

Sofroniew, M.V., Howe, C.L., and Mobley, W.C. (2001). Nerve growth factor signaling, neuroprotection, and neural repair. Annu Rev Neurosci 24, 1217–1281.

Spillane, M., Ketschek, A., Donnelly, C.J., Pacheco, A., Twiss, J.L., and Gallo, G. (2012). Nerve growth factor-induced formation of axonal filopodia and collateral branches involves the intra-axonal synthesis of regulators of the actin-nucleating Arp2/3 complex. J Neurosci 32, 17671–17689.

Stephens, R.M., Loeb, D.M., Copeland, T.D., Pawson, T., Greene, L.A., and Kaplan, D.R. (1994). Trk receptors use redundant signal transduction pathways involving SHC and PLC-gamma 1 to mediate NGF responses. Neuron 12, 691–705.

Summers, D.W., Frey, E., Walker, L.J., Milbrandt, J., and DiAntonio, A. (2020). DLK Activation Synergizes with Mitochondrial Dysfunction to Downregulate Axon Survival Factors and Promote SARM1-Dependent Axon Degeneration. Mol Neurobiol 57, 1146–1158.

Summers, D.W., Milbrandt, J., and DiAntonio, A. (2018). Palmitoylation enables MAPK-dependent proteostasis of axon survival factors. Proc Natl Acad Sci U S A 115, E8746–E8754.

Thomas, S.M., DeMarco, M., D’Arcangelo, G., Halegoua, S., and Brugge, J.S. (1992). Ras is essential for nerve growth factor- and phorbol ester-induced tyrosine phosphorylation of MAP kinases. Cell 68, 1031–1040.

Thornburg-Suresh, E.J.C., Richardson, J.E., and Summers, D.W. (2023). The Stathmin-2 membrane-targeting domain is required for axon protection and regulated degradation by DLK signaling. J Biol Chem 299, 104861.

Vohra, B.P., Sasaki, Y., Miller, B.R., Chang, J., DiAntonio, A., and Milbrandt, J. (2010). Amyloid precursor protein cleavage-dependent and -independent axonal degeneration programs share a common nicotinamide mononucleotide adenylyltransferase 1-sensitive pathway. J Neurosci 30, 13729–13738.

Walker, L.J., Summers, D.W., Sasaki, Y., Brace, E.J., Milbrandt, J., and DiAntonio, A. (2017). MAPK signaling promotes axonal degeneration by speeding the turnover of the axonal maintenance factor NMNAT2. Elife 6.

Wang, J., Shan, C., Cao, W., Zhang, C., Teng, J., and Chen, J. (2013). SCG10 promotes non-amyloidogenic processing of amyloid precursor protein by facilitating its trafficking to the cell surface. Human Molecular Genetics 22, 4888–4900.

Wang, J.T., Medress, Z.A., and Barres, B.A. (2012). Axon degeneration: Molecular mechanisms of a self-destruction pathway. Journal of Cell Biology 196, 7–18.

Wise, B.L., Seidel, M.F., and Lane, N.E. (2021). The evolution of nerve growth factor inhibition in clinical medicine. Nat Rev Rheumatol 17, 34–46.

Wood, K.W., Sarnecki, C., Roberts, T.M., and Blenis, J. (1992). ras mediates nerve growth factor receptor modulation of three signal-transducing protein kinases: MAP kinase, Raf-1, and RSK. Cell 68, 1041–1050.

Yamashita, N., and Kuruvilla, R. (2016). Neurotrophin signaling endosomes: biogenesis, regulation, and functions. Curr Opin Neurobiol 39, 139–145.

Yang, J., Weimer, R.M., Kallop, D., Olsen, O., Wu, Z., Renier, N., Uryu, K., and Tessier-Lavigne, M. (2013). Regulation of axon degeneration after injury and in development by the endogenous calpain inhibitor calpastatin. Neuron 80, 1175–1189.

Yao, R., and Cooper, G.M. (1995). Requirement for phosphatidylinositol-3 kinase in the prevention of apoptosis by nerve growth factor. Science 267, 2003–2006.

Yaron, A., and Schuldiner, O. (2016). Common and Divergent Mechanisms in Developmental Neuronal Remodeling and Dying Back Neurodegeneration. Curr Biol 26, R628–R639.

Zang, S., Ali, Y.O., Ruan, K., and Zhai, R.G. (2013). Nicotinamide mononucleotide adenylyltransferase maintains active zone structure by stabilizing Bruchpilot. EMBO Rep 14, 87–94.

